# *Cellos*: High-throughput deconvolution of 3D organoid dynamics at cellular resolution for cancer pharmacology

**DOI:** 10.1101/2023.03.03.531019

**Authors:** Patience Mukashyaka, Pooja Kumar, David J. Mellert, Shadae Nicholas, Javad Noorbakhsh, Mattia Brugiolo, Olga Anczukow, Edison T. Liu, Jeffrey H. Chuang

## Abstract

Three-dimensional (3D) culture models, such as organoids, are flexible systems to interrogate cellular growth and morphology, multicellular spatial architecture, and cell interactions in response to drug treatment. However, new computational methods to segment and analyze 3D models at cellular resolution with sufficiently high throughput are needed to realize these possibilities. Here we report *Cellos* (Cell and Organoid Segmentation), an accurate, high throughput image analysis pipeline for 3D organoid and nuclear segmentation analysis. *Cellos* segments organoids in 3D using classical algorithms and segments nuclei using a Stardist-3D convolutional neural network which we trained on a manually annotated dataset of 3,862 cells from 36 organoids confocally imaged at 5 μm z-resolution. To evaluate the capabilities of *Cellos* we then analyzed 74,450 organoids with 1.65 million cells, from multiple experiments on triple negative breast cancer organoids containing clonal mixtures with complex cisplatin sensitivities. *Cellos* was able to accurately distinguish ratios of distinct fluorescently labelled cell populations in organoids, with <3% deviation from the seeding ratios in each well and was effective for both fluorescently labelled nuclei and independent DAPI stained datasets. *Cellos* was able to recapitulate traditional luminescence-based drug response quantifications by analyzing 3D images, including parallel analysis of multiple cancer clones in the same well. Moreover, *Cellos* was able to identify organoid and nuclear morphology feature changes associated with treatment. Finally, *Cellos* enables 3D analysis of cell spatial relationships, which we used to detect ecological affinity between cancer cells beyond what arises from local cell division or organoid composition. *Cellos* provides powerful tools to perform high throughput analysis for pharmacological testing and biological investigation of organoids based on 3D imaging.

## Introduction

The selection of an optimal disease model is critical to the effective development and evaluation of cancer therapeutics. 2D monolayer cell cultures have been used extensively but have notable limitations. For example, although they can model cell autonomous characteristics, they lack multicellular aspects of the *in vivo* environment^1^ that may affect therapeutic performance, such as contributions of stromal components, spatial architecture, and cell polarity. Moreover, adaptation of cancer cells to adherent conditions on artificial substrates often perturb cellular characteristics. For such reasons, 3D cell culture models have emerged as a high-scale *in vitro* model platform for anticancer therapeutic discovery and development^2,3,4,5,6^. 3D profiling can enable broad new analysis possibilities because, unlike 2D profiling, it captures true cell spatial relationships and morphology, as well as potential cell-cell interactions. However, to realize these possibilities for high throughput treatment testing, improved 3D organoid data analysis is a key need^7^.

Organoid culture analysis is typically performed through image capture of organoids grown in multi-well plates, but available methods have limitations. Luminescence assays that are commonly used are cell-destructive methods that aggregate cell growth information over an entire well, whereas analysis of individual organoids and their constituent cells would be more informative. Current methods to identify organoids have focused on 2D segmentation, e.g. based on planar fluorescent quantification of live/dead cell stains and organoid areas^8,9,10,11,12,13,14^. Yet 2D analysis poorly approximates the complexity of 3D spatial relationships for cells and organoids^15^, which are likely to vary by cell type and growth conditions.

Moreover, prior computational approaches to analyze organoids in 3D have focused on qualitative visualization^7^ rather than cellular quantification, and current methods for 3D cell segmentation have been limited to specialized contexts. For example, Boutin et al.^16^ developed a 3D spheroid and nuclei segmentation approach for optically cleared images of one single spheroid in a well. More recently, Beghin et al. developed a segmentation technique for the Jewell system again with one organoid per well^17^. A more typical cancer treatment assay interrogates large numbers of organoids within each well of a plate. Development of a flexible 3D approach able to identify and quantify individual cells, as well as their morphologies, in high throughput could be of significant value to the field.

We demonstrate a novel computational method, *Cellos* (<u>Cell</u> and <u>O</u>rganoid <u>S</u>egmentation), to address this challenge. To our knowledge, *Cellos* is the first pipeline able to perform high-throughput volumetric 3D segmentation and morphological quantification of organoids and their cells, agnostic to the cell culture platform. *Cellos* has two stages. At first, organoids are segmented, and their volume, solidity, and other morphological characteristics are computed. In the second, nuclei are segmented in each organoid using a 3D convolutional neural network trained on an extensively curated new dataset. This enables analysis of characteristics including cell densities per well, clonal population frequencies, nuclear morphologies, and potential cell-cell interactions. We demonstrate the utility of *Cellos* by quantifying the complex three-dimensional responses of Triple Negative Breast Cancer (TNBC) organoids and their subclonal populations during platinum-based treatment *in vitro*.

## Results

### Description of the cancer cellular system

As a case study for 3D image analysis, we generated organoids from primary cell lines derived from a TNBC Patient Delivered Xenograft (PDX) model TM00099^18^. We used these cells because they have not been adapted to 2D cell culture conditions and retain clonal heterogeneity, and thus resemble the clinical *in vivo* situation. This TNBC model is *BRCA1* deficient and was initially sensitive to cisplatin. We have previously reported^18^ that this PDX tumor consists of two major subclones with differential cisplatin sensitivity. We isolated and established single cell-derived clonal lines, a cisplatin-resistant clone A50 and relatively sensitive clone B. We performed IC_50_ measurements for each of these clones individually grown as 3D organoids using a standard cell destructive luminescence assay, which assesses the number of viable cells from ATP content after cellular disruption (see methods for details). The A50 clone had an IC_50_ of 10.51 μM (95% CI= 8.71 μM - 12.44μM) and the relatively sensitive B clone had an IC_50_ of 2.94 μM (95% CI=2.61μM - 3.35μM) (Fig.1a). However, at high concentrations of cisplatin (>IC_80_), subclone B also exhibited a “dormancy” phenotype with elevated cell survival over A50. We took advantage of these clones to test the effectiveness of *Cellos* to quantify mixed populations with complex sensitivity profiles. We used nuclear localization signal (NLS)-conjugated EGFP or mCherry to stably label A50 and B clones and mixed them in two different ways: *homogeneously mixed organoids*, defined to be organoids made up of two genetically identical but differently labeled subclones (A50-EGFP with A50-mCherry, or B-EGFP with B-mCherry); and *heterogeneously mixed organoids*, defined to be organoids made up of two different subclones (A50-EGFP with B-mCherry, or A50-mCherry with B-EGFP). Organoids were imaged using the PerkinElmer Opera Phenix high-content screening system (Extended data fig. 1).

**Figure 1.**
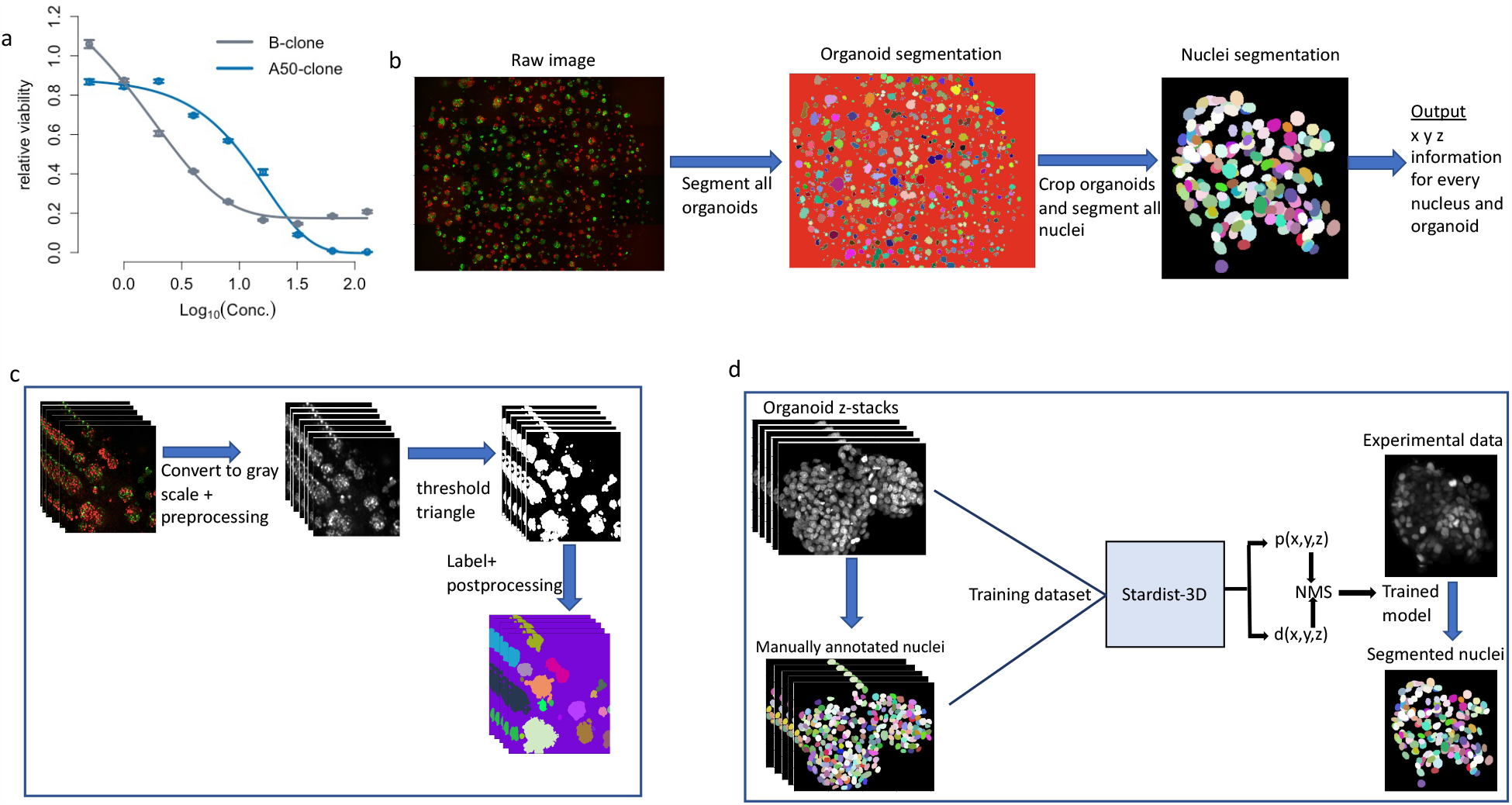
Outline of *Cellos* pipeline and cellular system. **a**. Cisplatin IC_50_ curves for two clones A50 (blue line) and B (gray line) from 3D homogeneously mixed organoids using cell-destructive luminescence readout. **b**. *Cellos*: Two-stage pipeline for 3D organoids and nuclei segmentation. **c**. Steps for 3D organoid segmentation. The inputs are 3D z-stack images, and the outputs are the segmented and labeled organoids. **d**. Steps for nuclei segmentation. A Stardist-3D with Resnet backbone model^22^ is trained using the training dataset. The trained model is then applied to experimental data with individual segmented organoids as input, and segmented and labelled nuclei as outputs.

### *Cellos* pipeline overview

The pipeline consists of two parts: organoid segmentation and nuclei segmentation (Fig.1b).

#### Organoid segmentation

To segment the organoids, first, we convert the fluorescent image to grayscale and preprocess to remove debris and noise (see methods for details). Next, we use the Triangle method for histogram thresholding^19^ to create a binary image separating the organoids from the background. We then use scikit image^20^ to uniquely label all organoids, remove small objects (Fig.1c), and generate a table of measurements for the remaining organoids (3D bounding box, volume, mean intensity, solidity, etc.). All steps are performed on small fields defined by the imaging platform used and then stitched together before the labeling step. A csv file that contains measurements of all organoids is generated for each well in the plate, allowing parallel processing of individual wells from the same plate.

#### Nuclei segmentation

For each well, stitched z-stack images and the measurement file for each individual organoid are used as inputs. Segmentation of cells within the organoids by classical methods is challenging due to the high density of cells in organoids. Therefore, we developed a convolutional neural network (CNN) for nuclei segmentation using the Stardist-3D^21^ model with a ResNet backbone^22^. To train the model, we generated a training dataset of manually annotated 3,862 nuclei in 3D from 36 TNBC organoids with a range of 8-440 cell nuclei. The organoids consisted of EGFP, mCherry or Hoechst labelled cells and 24 of the 36 organoids were imaged after exposure to a range of cisplatin concentrations. We then applied the trained model to experimental data (Fig.1d). We also used scikit-image to generate measurements (centroid, volume, mean intensity, solidity, etc.) for every nucleus. Our trained CNN is publicly available on the *Cellos* GitHub.

### *Cellos* organoid segmentation is accurate

To evaluate the effectiveness of *Cellos* for segmenting organoids, we applied it to heterogeneously mixed organoids (A50-EGFP with B-mCherry) in several wells. These wells were treated with cisplatin concentrations ranging from 0-128 μM (see methods dataset-1 for details). Organoid segmentation appeared accurate in the absence (Fig.2a, top panel) and presence (Fig.2a, middle and bottom panels) of cisplatin. To quantify this, we manually counted true positives, false positives, and false negatives for 321 segmented organoids in wells treated with different concentrations of cisplatin (16-128 μM). We focused on high cisplatin doses because they are more difficult to accurately segment: their organoid shapes are more diffuse, and the images have higher cell debris and noise. We observed precision, recall, and F1 scores of 96.07, 83.80, and 89.52, respectively, where the F1 score is the harmonic mean of recall and precision.

**Figure 2:**
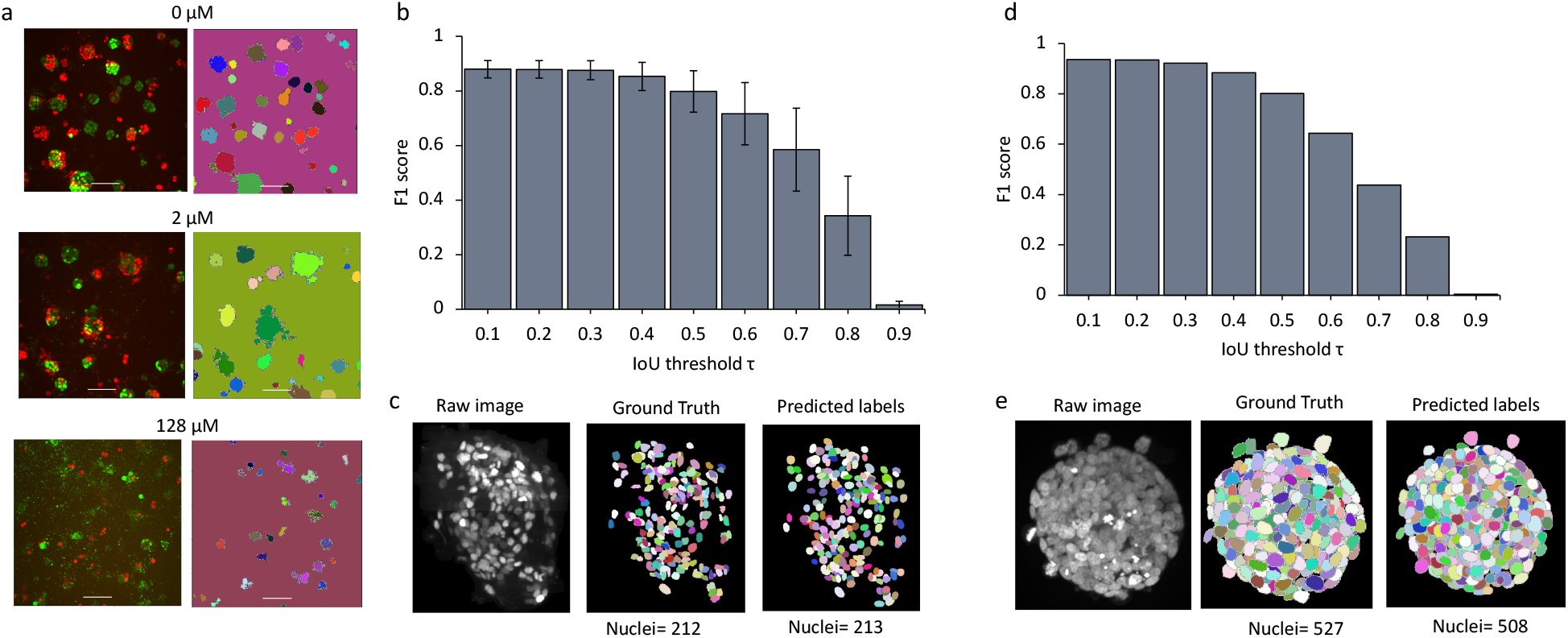
Evaluation of organoid and nuclei segmentation. **a**. For each pair of images, the left shows fluorescence z-axis maximum projections with A50 cells labeled with EGFP and B cells labeled with mCherry. The right panel shows organoids segmented by *Cellos*. Individual organoids are in distinct colors. Organoids from untreated, 2 μM, and 128 μM cisplatin wells are shown. **b**. F1 score for organoid identification vs. IoU threshold. Mean and standard deviation across six cross-validations is shown. **c**. Example EGFP-labeled organoid image (left panel) with manually annotated ground truth nuclei annotations (middle panel) and *Cellos* predicted labels (right panel) respectively. Images are z-axis maximum projections. **d**. F1 score of *Cellos* nuclei segmentation in spheroids of breast carcinoma from Boutin et al.^15^ vs. IoU threshold. **e**. Example DAPI stained image (left panel) from Boutin et al.^15^ with ground truth (middle panel) and *Cellos* predictions (right panel), respectively.

### *Cellos* nuclei segmentation is accurate and robust on independent data

To evaluate *Cellos* for nuclear segmentation, we first used internal cross-validation. We performed six-fold cross-validation (30 and 6 organoids for training and validation respectively) and computed F1 score versus intersection over union (IoU) thresholds. IoU is the spatial overlap between the ground truth and the predicted nuclear region; the bigger the IoU, the greater the overlap. We observed high F1 scores extending to relatively large IoU thresholds, example F1 score of 0.853±0.052 at IoU of 0.4 (Fig.2b), indicating the quality of the match between predicted nuclei and ground truth. Visual inspection supported the consistency in ground truth and predicted labels, and the total number of nuclei identified were similar in the predictions and ground truth (Fig.2c). These results show the accuracy of *Cellos* to segment nuclei, despite the challenge of high anisotropy in z resolution relative to x and y^21^.

We next tested whether *Cellos* could segment nuclei from organoids in external datasets generated independently from our group. We used data from Boutin, et al.^15^, who manually annotated cells in three optically cleared spheroids grown and assayed in a manner distinct from our platform. The spheroids were from a breast carcinoma cell line T47D grown in 384-well round-bottom Ultra-Low Attachment plates for 3 days. They were stained with DAPI and imaged using a 20x water objective magnification with 5 μm z-step size. Despite these system and fluorescence marker differences, *Cellos* accurately segmented the nuclei (1585 nuclei) in their spheroids with F1=0.885 at IoU of 0.4 (Fig.2d), with an example spheroid shown in (Fig.2e). We compared Cellos to the 3D classical nuclei segmentation protocol that Boutin et al. developed for their spheroids. *Cellos* showed superior specificity of 93.38% and F1= 0.94 compared to the Boutin et al. method with 75.76% for specificity and F1=0.76.

### *Cellos* precisely detects distinct proportions of labelled cells in mixed organoids

The ability to quantify multiple cell types simultaneously in an organoid can enable studies of cancer population dynamics. To evaluate the efficacy of *Cellos* for this task, we analyzed homogeneously mixed organoids (A50-EGFP with A50-mCherry) at four different seeding ratios of EGFP-20%, EGFP-40%, EGFP-60%, and EGFP-80%. These organoids were imaged at day0, allowed to grow, and imaged again 4 days later (day4) (see methods dataset-2 for details). We performed similar experiments for homogeneously mixed organoids of B-EGFP with B-mCherry.

We observed that the EGFP fluorescence increased with the EGFP seeding ratio in organoids consisting of A50-EGFP with A50-mCherry (Fig.3a). To evaluate this quantitatively, we used *Cellos* to calculate the total number of EGFP and mCherry cells in each well and their corresponding ratios. Indeed, across all wells imaged at day0, we detected ratios close to the expected EGFP:mCherry ratios, with average absolute difference of 2.986% (Fig.3b). Ratios were also stable from day0 to day4, with mean difference of 2.852%. We calculated the deviation for each set of triplicate wells, and the average standard deviation across seeding conditions was 0.808% (Fig.3b). We repeated these experiments for organoids generated from mixtures of B-EGFP and B-mCherry and found similar results (Fig.3b). We did not detect any bias towards either fluorescent label or clone type (extended data table 1).

**Figure 3.**
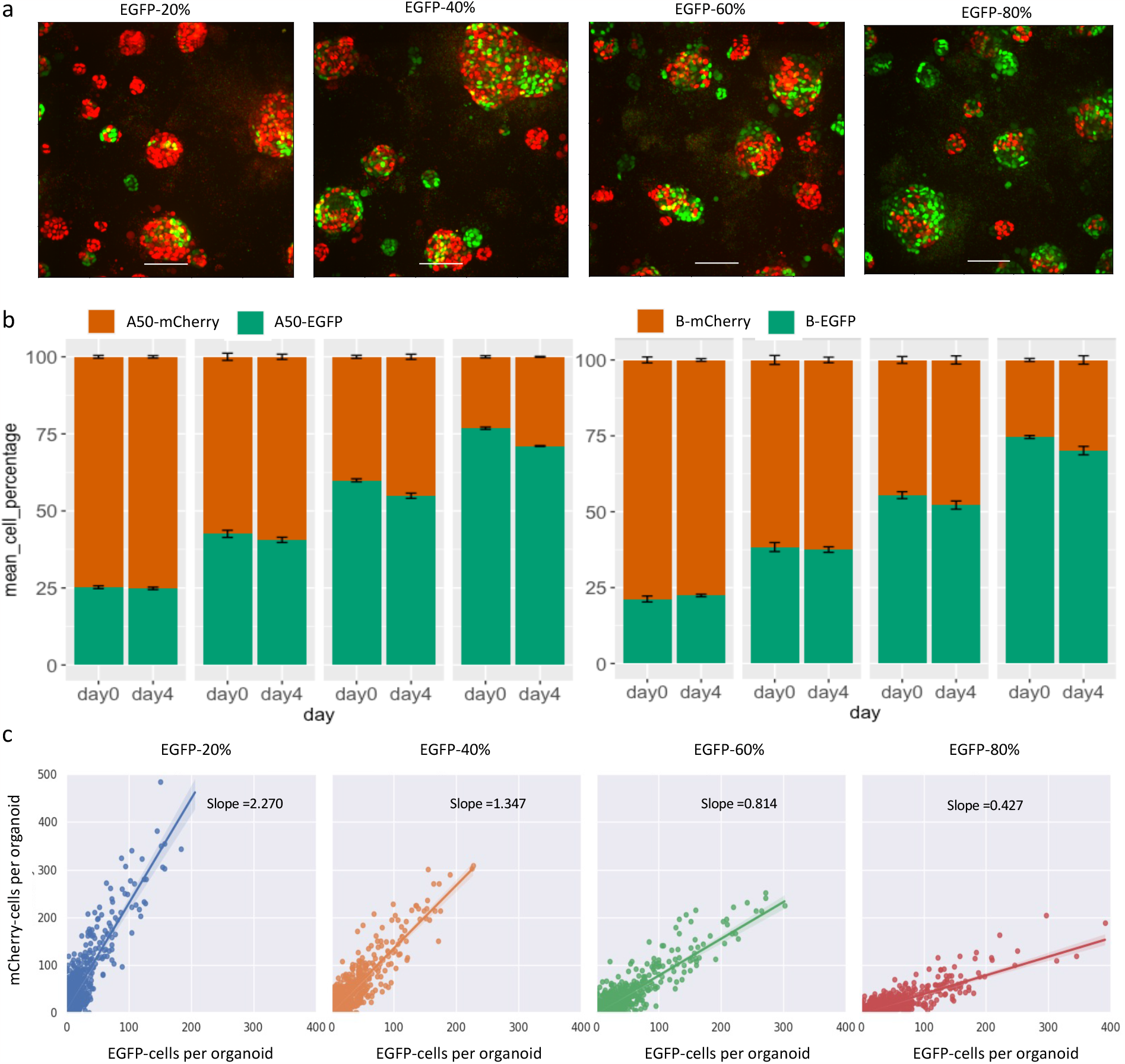
*Cellos* distinguishes proportions of co-cultured cells. **a**. Representative z-axis maximum projection images of homogeneously mixed organoids generated with seeding percentages of 20%, 40%, 60% and 80% A50-EGFP, respectively, with the remaining cells being A50-mCherry. Images are at day4. Scale bar represents 100 μm. **b**. Stacked bar plot showing the percentage of EGFP and mCherry cells detected by *Cellos* at day0 and day4 for each of the seeding conditions from Fig.3a (left panel). Analogous experiments for B-EGFP+B-mCherry mixed organoids (right panel). Error bars indicate standard deviation calculated from three replicate wells for each condition. **c**. Number of cells labelled with EGFP vs. mCherry detected in each homogeneously mixed A50 organoid. Each dot depicts an organoid. Seeding conditions of EGFP-20% (blue), EGFP-40% (orange), EGFP-60% (green) and EGFP-80% (red) are shown from left to right. The slope of the fitted linear regression is noted.

Next, we evaluated how these ratios varied within individual organoids across the different seeding conditions. Using *Cellos*, we quantified the number of EGFP and mCherry cells in each organoid in each well (total 561,722 nuclei and 23,258 organoids) and plotted these values across organoids (Fig.3c) for each seeding condition. As expected, the slopes of the regression line between number of EGFP versus mCherry cells changed monotonically with seeding ratio. Specifically, we observed slopes of 2.270, 1.347, 0.814, 0.427 for the wells with seeding conditions of EGFP-20%, 40%, 60%, and 80%, respectively for A50 (Fig.3c). Observations were similar for B (Extended data fig.2). This supports that *Cellos* is able to discern cellular ratios across different organoid sizes.

### *Cellos* robustly quantifies treatment response of co-cultured clones

An important use of *Cellos* is to quantify treatment response profiles of organoids made up of multiple clones, a task which may be valuable for parallelizing response evaluations and detecting clonal interactions. The A50 and B clones from TM00099 tumors have distinctive drug responses to cisplatin (Fig.1a) making them well-suited to evaluating *Cellos* for this task. We generated heterogeneously mixed organoids consisting of A50-EGFP and B-mCherry cells and treated them with a range of (0-128 μM) cisplatin concentrations (see methods dataset-1) in triplicates. We first used *Cellos* to estimate cell density in each well. As expected, we saw a decrease in the cell density as cisplatin concentration increases (Fig.4a), confirming effectiveness of treatment. We then used clone-specific cell densities to determine IC_50_ for each clone. A50 was 2.7x more resistant to cisplatin than B: A50 IC_50_ = 2.87μM (95% confidence interval (CI) = 1.95μM – 4.54μM) and B IC_50_ = 1.06 μM (95% CI=0.87μM - 1.35μM), respectively (Fig.4b). At higher doses (∼IC_80_), and as expected, we also observed a relatively higher viability of B than A50. These observations were consistent with the standard luminescence assays (Fig.1a), indicating that *Cellos* can recapitulate those findings by counting individual cells of specific cell types, but in a non-destructive and higher (cell) resolution manner.

**Figure 4.**
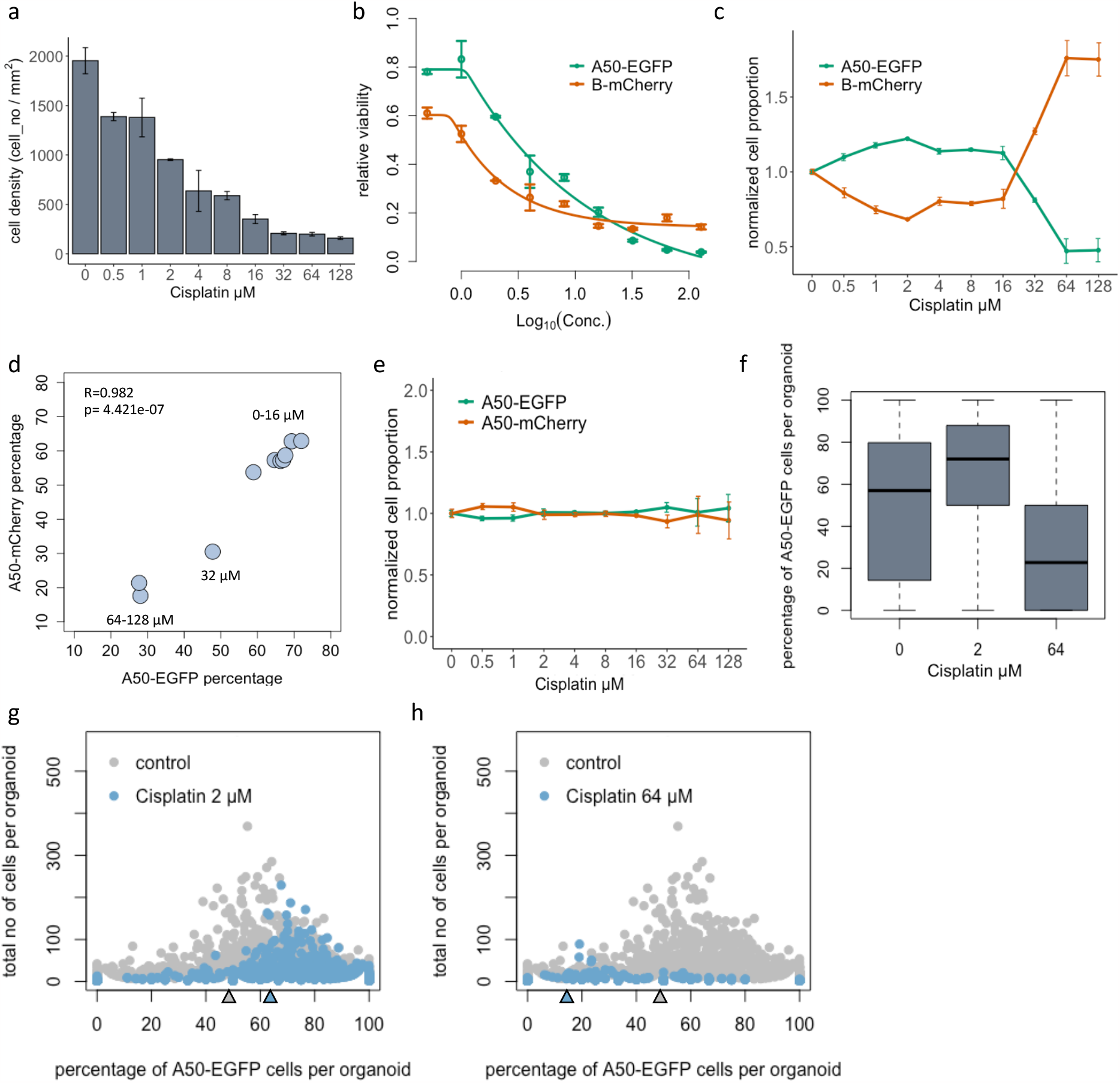
*Cellos* quantification of treatment response for organoids consisting of co-cultured clones. **a**. Total cell density (A50 and B) after exposure to cisplatin (0-128 μM) for 4 days. Mean and standard deviation across three replicate wells per condition are shown. **b**. IC_50_ curves for A50-EGFP and B-mCherry clones when co-cultured in heterogeneously mixed organoids, with standard deviation across triplicates shown for each condition. **c**. Normalized cell proportions vs. cisplatin treatment concentration range, for A50-EGFP and B-mCherry clones cultured as mixed organoids. **d**. Correlation of A50 clonal percentages post cisplatin treatment, when labeled with either nuclear EGFP or mCherry and mixed to form organoids with B clones labelled with the contrasting fluorescent channel. **e**. Normalized cell proportions vs cisplatin treatment concentration range, for A50-EGFP and A50-mCherry clones co-cultured as mixed organoids. **f**. Distribution of percentage of A50-EGFP cells per organoid when mixed with B-mCherry cells after 0, 2 and 64 μM cisplatin exposure. Triplicate wells are merged. **g**. Percent A50-EGFP cells per organoid versus total cells. Each dot represents an organoid. Gray = untreated. Left panel: Blue = 2 μM cisplatin. Right panel: Blue = 64 μM cisplatin. Black arrow indicates median for the control. Red arrow indicates median for the treatment.

To further validate these results, we analyzed the ratio of A50 and B cells in the cisplatin-treated wells containing heterogeneously mixed organoids. For each drug condition, the ratio of A50-EGFP to B-mCherry cells per well was computed (Extended data fig.3a) and normalized for the ratio of the respective clones in untreated wells. We observed increasing A50-EGFP and decreasing B-mCherry normalized cell proportion with increasing concentrations of cisplatin, up to 16 μM (Fig.4c). This trend reversed at higher concentrations of 64 and 128 μM cisplatin, consistent with the luminescence and *Cellos* based cell density estimates. This confirms the presence of a dormancy phenotype in B.

To determine whether fluorescence labeling might have impacted cell viability or generated any imaging bias, we flipped the fluorescent labels and repeated the experiment, now generating and treating heterogenous organoids with A50-mCherry and B-EGFP cells (see methods, dataset-3). The flipped-label results were highly correlated with the originals (R=0.982, Fig.4d, see also extended data fig.3b), confirming that the observed treatment responses were determined by inherent clonal differences. We also analyzed homogeneously mixed organoids consisting of A50-EGFP and A50-mCherry cells treated with cisplatin (see methods, dataset-4). As expected, the normalized proportions of EGFP and mCherry cells were stable across all cisplatin conditions (Fig.4e). Variations in IC_50_ and cell density curves across replicates were affected by variable spatial distribution of organoids relative to the imaged regions of each well. However, the ratio of EGFP to mCherry was highly consistent across replicates (Fig.4c) since they are independent of total cell numbers imaged per well.

At the individual organoid level, we observed similar clonal compositional shifts but with greater stochasticity. The median A50-EGFP percentage across organoids increased from 52 % to 72 % from 0 to 2 μM and decreased to 23 % at 64 μM, with variability amongst individual organoids (Fig.4f and g). In particular, for organoids with few cells we observed wider variation in the A50-EGFP percentage, as expected from their susceptibility to stochastic fluctuations. We also computed mean fluorescence intensities of the clones in each organoid, and these yielded findings similar to those from the cell counts (extended data fig.3c).

### *Cellos* can reveal organoid and nuclear morphology changes due to therapy

Organoid morphology may change in response to therapy and is commonly assessed to evaluate effects of drugs on organoids in qualitative image-based assays^23^. We therefore investigated whether *Cellos* could detect quantitative differences in morphology among organoids after cisplatin treatment. Multiple morphological features (volume, volume filled, volume convex, volume bbox, intensity mean, intensity max, intensity min, eccentricity, solidity, Euler number, and inertia tensor eigvals) for every organoid were computed using *Cellos*. Several features exhibited clear changes post treatment. We observed monotonic decreases in organoid volume (Fig.5a), solidity as a measure of cell packing (Fig.5b), and total cell number per organoid (Fig.5c) as drug concentration increased. These factors showed statistically significant differences (p<0.05) compared to control even at low cisplatin thresholds, e.g. solidity declined at 1 μM cisplatin, p= 4.13e-10, cell number also decreased at 1 μM Cisplatin, p=5.87e-05, and organoid volume was reduced at 4 μM cisplatin, p= 4.44e-08.

**Figure 5.**
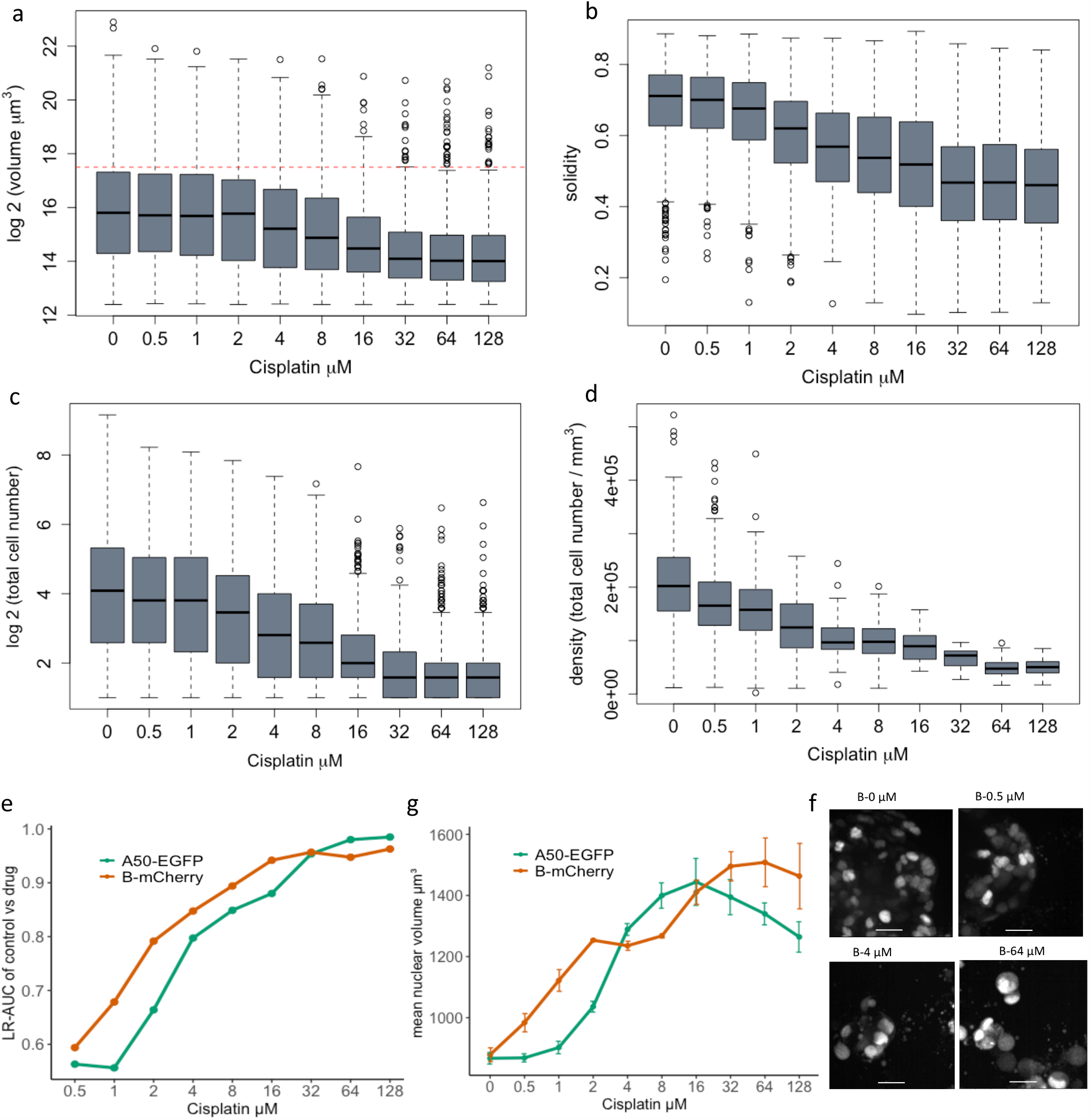
Changes in organoid and nuclear morphological features after cisplatin treatment. **a**. Organoid volume (μm^3^), **b**. Solidity and **c**. Cell number after cisplatin treatment for 4 days. **d**. Cell density within large (>1.85×10^5^ μm^3^) organoids after 4 days of cisplatin exposure. **e**. Logistic regression classifications based on nuclear morphology, for classification of control nuclei vs. cisplatin-treated nuclei. A50-EGFP and B-mCherry cells are analyzed separately. LR-AUC indicates area under the curve of the logistic regression classifier for the specified comparison. **f**. Nuclear volume of A50-EGFP and B-mCherry cells for a range of cisplatin concentrations. Mean values with standard deviation for triplicate wells are plotted. **g**. Representative images of B-mCherry cell nuclei at day 4 post cisplatin exposure. Scale bar represents 25 μm.

Despite the decrease in average volume at high cisplatin concentrations, there were still a number of remnant large organoids. 2.5 % of organoids in the 64 and 128 μM cisplatin wells had volume greater than 1.85×10^5^ μm^3^ (red dotted line in Fig.5a), the threshold for the largest quartile of untreated organoids. Such large organoids (>1.85×10^5^ μm^3^) exhibited different morphological characteristics in the untreated versus treated wells. For example, large organoids showed 4.25-fold lower cell density in the 64 μM wells compared to untreated organoids (Fig.5d). The treated large organoids also showed lower solidity, lower eccentricity (minor/major axis), and higher Euler number (measure of holes present) (Extended data fig. 4a-d). This suggests that the structure of the large organoids appears to be maintained despite a reduction in cellular content (as measured by lower cell density, solidity, and higher Euler number). Thus, *Cellos* is able to detect subtle but distinctive morphological changes at the organoid level as the result of escalating doses of chemotherapy.

We next studied whether *Cellos* could classify response based on nuclear morphology changes caused by cisplatin treatment. To do this, we trained binary logistic regression models to classify untreated cells versus treated cells for each concentration of cisplatin treatment. A total of 235,957 nuclei were used for this analysis. Separate sets of classifiers were trained for the A50-EGFP and B-mCherry clones. The inputs to the classifiers were *Cellos*-identified morphological features (volume, volume filled, volume convex, volume bbox, intensity mean, intensity max, intensity min, eccentricity, solidity, Euler number, and inertia tensor eigvals) for every nucleus. For each model, 80% of the dataset was used for training and the remaining 20% was used for testing. In addition, area under the ROC Curve (AUC) was used to evaluate the model. As expected, the logistic regression classifiers showed increasing AUC for cells exposed to higher cisplatin concentrations for both clones (Fig.5e). However, even at low drug concentrations, the two clones, A50 and B already started to show morphological differences with ∼0.6 AUC at 1 μM, to 0.7 AUC at 2 μM cisplatin. At higher cisplatin concentrations (>4 μM), the classifier gave AUCs of >0.8 and at the highest dose of cisplatin (128 μM) the AUC was >0.95 for the classification of untreated versus cisplatin treated A50 and B nuclei. This suggests that 3D nuclear morphology can be used as a quantitative surrogate for chemotherapeutic effect.

To verify that these results were not fluorescent label-specific, we repeated the classifier analyses using the reversed fluorescent-labelled nuclei (B-EGFP and A50-mCherry). AUCs were robust despite label flipping (extended data fig.5a-b), with the maximum AUC difference across the flips being only 0.0323, observed at 64 μM for B-mCherry vs. B-EGFP. This confirms that when a cell is exposed to cisplatin, changes in their nuclear morphology occur in a dose dependent manner, and that these changes are sufficient to build discriminative models regardless of the fluorescence marker used.

Nuclear volume and fluorescence intensity were the morphological features that provided the greatest contribution in the logistic regressions. Thus, we further analyzed these features individually. Fluorescence intensity decreased with cisplatin concentration for both clones (extended data fig.5c), possibly indicating that the cellular stress induced by drug exposure might reduce either label’s transcription or translation. Nuclear volume had a more complex behavior, initially increasing with increase in cisplatin concentrations for both A50 and B (Fig.5f), which was also apparent by visual inspection (B clone: Fig.5c, A50 clone: extended data fig.5d). Interestingly, the B clone showed a faster increase in nuclear volume compared to the A50 clone at lower doses of cisplatin, which is consistent with the logistic regression AUC values and with the greater sensitivity of B clones to cisplatin (Fig.5e). However, A50 cells showed a significantly higher decline in nuclear volume at concentrations >32 μM cisplatin than B clone. Decrease in nuclear sizes have been observed in cells underdoing apoptosis^24,25^. Thus, the early reduction in A50 nuclei volumes could indicate the initiation of apoptotic pathways which would correlate with the increased cell death in A50 clones at high cisplatin concentrations compared to the B clone that exhibits a persister-like behavior^26^. While this confirms that B cells survive better at high drug concentrations, *Cellos* also provides new information that these surviving cells are not normal and show cellular stress indicated by relatively larger nuclear volume. While the precise biological meaning of these morphological changes is not clear, the ability to have refined discernment of morphological alterations on a single cell level can motivate more detailed experimentation, especially around the biology of distinct cellular responses to different levels of drug exposure. In the case of nuclear volume, our observations potentially suggest that the nuclear effects at cytotoxic doses <32 μM are different from the nuclear effects after exposure to doses >32 μM cisplatin.

### *Cellos* reveals spatial relationships between cells in organoids

3D segmentation at cellular resolution allows for analysis of cell spatial relationships, which may reveal ecological interactions among clones. To interrogate such interactions, we analyzed how clones were organized within heterogeneously mixed TNBC organoids. For each clone in each organoid, we calculated the *localization score*, which quantifies how often a cell’s adjacent cells are of the same clonal type, within a specified cell number window size. This score is normalized for the clonal fraction in the organoid. The higher the localization score, the higher the co-localization of cells of the specified clone (see methods).

We observed that cells of a given clone tend to co-occur with others of the same clone, as demonstrated by a decrease in localization score with increasing window size. This was true both in the presence and absence of cisplatin treatment, and for both clones B (Fig.6a) and A50 (Extended data fig.6a). These results indicate that clones form small spatial clusters within the organoids. Co-localization was stronger in the organoids having only a small fraction of one clone, whether the minor population clone was B (Fig.6b) or A50 (extended data fig.6b). This effect was consistent even in the 10 cell windows (extended data fig.6 c, d). An example of a heterogeneously mixed organoid (consisting of 0.202% B clone fraction) with strong clone localization (localization score=3.1) is shown in Fig.6c. In organoids with equal fractions of the two clones, we observed cases both where clones are well mixed (Fig.6d, localization score=1.25) and where they form separate clusters within the organoid (Fig.6e, localization score=1.53).

**Figure 6:**
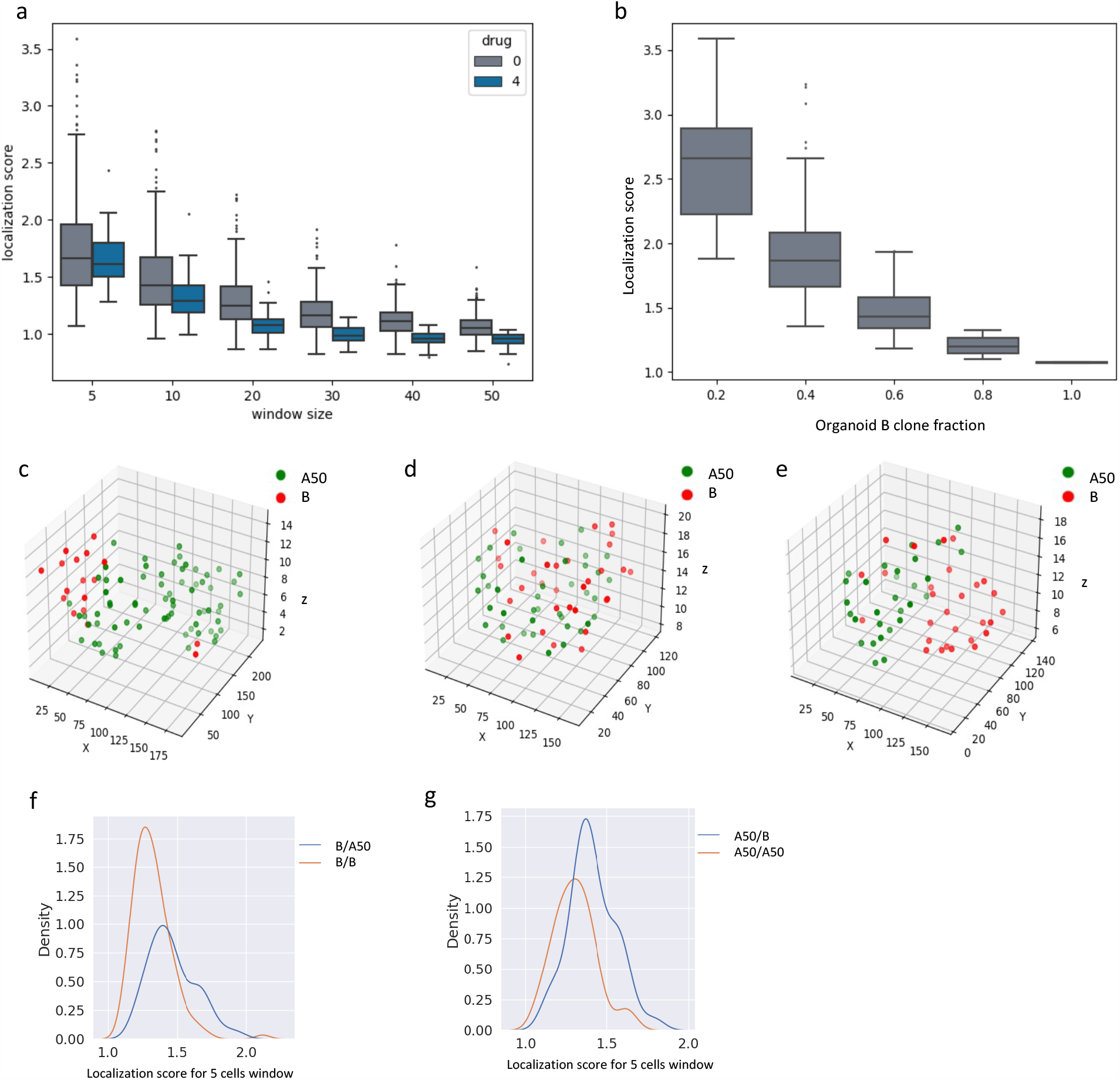
Investigation of cell-cell spatial relationships within organoids. **a**. Localization score vs. window size (5, 10, 20, 30, 40 and 50 cells) for B-mCherry cells within heterogeneously mixed organoids. Control and 4 μM cisplatin data are shown. **b**. Localization score for B-mCherry cells in organoids (5 cell window), stratified by B clone fraction in the organoid. Organoids are binned in increments of 0.2 for the B cell fraction. **c, d, e**. Spatial location of nuclei centroids in representative organoids plotted in 3D. **f**. Distribution of localization score of B clones (in 5-cell window) when mixed with A50 (blue line) or with alternately labeled B (orange line). **g**. Distribution of localization score of A50 clones when mixed with B (blue line) or with alternately labeled A50 (orange line).

Local cell division could generate spatial clusters of cells within organoids, but they could also arise from ecological affinity between cells of the same clone. Comparing homogenously mixed organoids to heterogeneously mixed organoids allows one to distinguish whether spatial clusters arise from cell division or ecological affinity. This is because homogeneously mixed organoids should demonstrate the cell division effect but not the ecological affinity effect. We observed significantly higher clone localization in heterogeneously mixed organoids compared to homogeneously mixed organoids (Fig.6 f, g. This was the case for comparison of homogeneous B organoids to B-A50 mixed organoids (p=1.5e-9), as well as for comparison of homogeneous A50 organoids to A50-B mixed organoids (p=5.2e-6). Thus, these clones have greater ecological affinity for cells of their same type. Note that to avoid the effect of clonal fraction on localization score, we restricted these analyses to organoids with comparable clonal fractions, within a range of 0.4-0.6.

## Discussion

We have presented *Cellos*, a high-throughput pipeline for segmenting and analyzing organoids at the cellular level. Novel advantages of *Cellos* include its abilities to accurately segment organoids and cells in 3D from distinct imaging modalities; to robustly determine cell ratios in wells; to generate IC_50_ curves based on cell segmentations within images; and to identify organoid and nuclear morphological features associated with treatment response. Moreover, *Cellos* enables 3D analysis of cell spatial relationships, which are valuable to the assessment of tumor microenvironmental interactions.

The high-throughput capacity of *Cellos* is crucial to its value for drug or gene knockout screening analysis. To clarify its scalability, 74,450 organoids and 1.65 million cells were analyzed for the experiments presented here. As *Cellos* has the ability to process multiple wells of a plate at the same time, the time used to investigate 60 wells was similar to the one used for one well. Imaging speed and size of the data in 3D are also important practical considerations for large scale image-based assays. To minimize imaging time, we used coarser z-sampling intervals of 5 μm while still achieving accurate segmentation, requiring far less data. Prior studies have used z-stack intervals of 0.122 μm to study C. elegans^27^ and 1.2 μm intervals for cleared spheroids^28^. Axial sampling of 5 μm was chosen because our initial studies indicated that resolutions finer than 5 μm did not appreciably improve segmentation accuracy. Additionally, we limited the number of z-slices to 100 and imaged the same z-range for all wells in a plate, which restricted identification of organoids to certain regions of the well. Because organoid distribution within wells can be variable, this led to differences in cell counts across replicate wells. To address this problem, *Cellos* includes a customized method for distinguishing image areas that contain organoids in focus within the acquired image of every well (extended data fig.7a-c). This area estimate can then be utilized to determine organoid and cell densities, which allow for a more robust comparison between wells. However, quantifications based on clonal ratios within a well are less sensitive to this issue, since ratios account for absolute cell numbers. Furthermore, to decrease storage, images generated by *Cellos* were saved as zarr arrays^29^, which then required ∼12GB to store an image of one well (with a size of 3×101×5080×5080 pixels). Altogether, when pre-processing and morphological feature characterization are included, it took on average ∼1.9 hours with CPU efficiency of 91.2% and ∼100 GB of computational memory to segment 550 organoids, and ∼1.4 hours with CPU efficiency of 82.31% and 6.86 GB of computational memory to segment 4837 nuclei.

A major challenge in cancer therapeutics is deciphering the effects of multicellular interactions. For example, the tumor microenvironment contains diverse tumor clones, immune cells, endothelial cells, and tumor-associated fibroblasts, which may each provide new target pathways for drug development^30,31,32^. Non organoid approaches to evaluate how drugs interact with the tumor microenvironment have included measurements in animal models, tissue explants, or observations in clinical samples^33^, all of which are challenging to scale. In contrast, organoid technologies allow interrogation of drug and cellular combinatorics based on high numbers of independent multicellular structures. *Cellos* enables interpretation of these parallel measurements by individually quantifying cells and organoids to elucidate chemical biology and pharmacology effects. *Cellos* is effective in 3D cell segmentation despite complexities such as the difficulty of identifying cell boundaries in packed organoids, as well as inherently large differences in cellular morphology of different cell types which can change further post drug treatment. It is worth noting that our use of nuclear labels or DNA counterstains was valuable in enabling *Cellos* to segment cells robustly enough for these tasks.

To clarify the phenotypic importance of 3D quantification, we investigated the dynamics of triple negative breast cancer organoids having complex multiclonal cisplatin sensitivity. Using *Cellos*, we could accurately recapitulate the IC_50_ curves of well-based luminescence assays, including subtle non-monotonic changes in relative drug sensitivity. Moreover, the morphological characterizations enabled new associations that have not been previously possible. For example, we were able to quantify how nuclear volume changes with cisplatin, and how volume tracks with viability changes observed in IC_50_ curves. While we have focused on how intuitive features such as volume and fluorescence intensity contribute to logistic classifiers, using more general image analysis algorithms, such as convolutional neural network^34,35^may provide an even greater potential for classification of the biological states of organoids and cells.

Another promising direction enabled by *Cellos* is the quantification of cell-cell spatial relationships - an inherently three-dimensional problem. We were able to distinguish the affinity of a cancer cell clone to localize with other cells of the same clone, from the effects of local cell division. An interesting future direction would be to study if such affinities can modulate treatment response. In any case, the detection and quantification of ecological affinities in model systems will be valuable for understanding the interactions within tissue microenvironments. Such analyses can clarify mechanisms in which cell juxtapositions result from factors including cell division, motility, cytokine signaling, and cell-cell ligand-receptor interactions.

In conclusion, we report the first high-throughput-compatible pipeline for 3D organoid and nuclei segmentation, as well as morphological quantification. *Cellos* opens new opportunities for organoid experimental design and the elucidation of multicellular phenotypes from imaging.

## Materials and Methods

### Clonal line establishment and nuclear fluorescent labelling

Primary cell cultures were derived from TM00099 PDX tumor fragments as previously described^18^. Briefly, tumor fragments were disassociated in collagenase (1-2 hours, 37°C), and harvested cells were washed and plated on irradiated 3T3-J2 feeder cells and maintained in 37°C and 7.5% CO_2_ in culture medium as described by Liu et al.^37^ After *in vitro* expansion of the primary cell lines, single human cells labelled with Anti-Human HLA-ABC APC (eBiosciences) were isolated using flow cytometry and further expanded to establish clonal lines. Two clonal lines defined as A50 and B were then stably transduced with lentivirus expressing nuclear EGFP or mCherry. pTRIP-SFFV-EGFP-NLS (Addgene) was used for the lentiviral nuclear EGFP construct. mCherry sequence was subcloned from pLenti6-H2B-mCherry (Addgene) to replace the EGFP sequence thus generating a pTRIP-SFFV-mCherry-NLS construct.

The two lentiviral plasmids were packaged into lentivirus by transfecting 5 μg of each plasmid with psPAX2 (Addgene) and pMD2.G (Addgene) in HEK293T (ATCC) cells with Lipofacamine 2000 (Invitrogen). Lentivirus collected after transfection was concentrated using 3P Lenti-X™ Concentrator (Clontech Labs) using manufacturers protocol. Clonal cell lines were infected with the lentivirus and selected for by sorting for nuclear EGFP or mCherry cells respectively. All clonal lines were authenticated by PCR amplification of clonal line specific structural variations and routinely tested for Mycoplasma contamination using the MycoAlert PLUS Mycoplasma Detection Kit (Lonza).

### 3D cell culture assays and imaging

For 3D culture assays, fluorescence-activated cell sorting (FACS)-sorted EGFP or mCherry cells were seeded at a density of 30,000 cells per well in triplicate in 96 well plates (PerkinElmer) coated with 35 μl Matrigel Growth Factor Reduced (BD Biosciences) on day -3. Cells were cultured in 37°C and 7.5% CO_2_ for three days for organoids to form. On day 0, media was replaced, and organoids were treated with 9 doses of cisplatin (Selleck Chemicals) in two-fold serial dilutions in the range of 0.5 to 128 μM. After 96 hours of drug exposure, on day 4, CellTiter-Glo® 3D Cell Viability Assay (Promega) was used for luminescence-based assay readouts using manufacturer’s protocols.

For image-based readouts, plates were imaged at day 0 or day 4 using the Opera-Phenix High-Content Screening System (PerkinElmer). 25 fields per well, were imaged using a 20x water objective. In each field, 101 z-stacks at 5 μm separation were imaged in three channels - brightfield, EGFP and mCherry. Cells were maintained at 37°C and 7.5% CO_2_ during the imaging. For training set experiments, nuclei were counterstained with a final concentration of 1 μg/ml Hoechst 33342 Solution (Thermo Scientific™) for 30 minutes at 37°C before imaging.

For generation of heterogeneously mixed organoids, A50-EGFP and B-mCherry cells (dataset-1) or A50-mCherry and B-EGFP (dataset-3) cells were mixed in 50:50 ratios and treated with drug as described above and imaged at day4 post drug exposure. For generation of dataset-2, homogeneously mixed organoids consisting of A50-EGFP and A50-mCherry (or B-EGFP and B-mCherry cells) were generated by mixing them in varying rations of EGFP:mCherry namely – 20:80, 40:60, 60:40 or 80:20 respectively. These organoids were imaged at day 0 and day 4. For datasets 4 and 5, homogeneously mixed organoids consisting of equal proportions of A50-EGFP and A50-mCherry (dataset-4) or B-EGFP and B-mCherry cells (dataset-5) were generated and treated with cisplatin and imaged as described above.

### Analysis protocol overview

We developed a customized pipeline combining python and shell scripts to analyze 3D organoids both at the organoid and cellular level. The customized pipeline is written to process an entire multi-well plate. Multiple wells can be processed at the same time on a high-performance computing cluster by taking advantage of multiple *central processing unit* (CPU) cores. The main steps for the pipeline are: 1. Exporting and organizing image data, 2. 3D segmentation of individual organoids, 3. Cropping individual organoids and 3D segmentation of nuclei in each organoid, 4. Computing morphological features and saving output information, 5. Calculating area of imaged well with organoids, 6. Post segmentation analysis like: IC50 estimates and cells co-localization analysis. The pipeline code is available on GitHub at https://github.com/TheJacksonLaboratory/Cellos.

### Exporting and organizing image data

The image data were exported from the Opera Phenix high content screening confocal microscope using the Harmony High-Content Imaging and Analysis Software (PerkinElmer). The resulting folder contains subfolders with tiff files (Images) and xml file (metadata). Each tiff file is a single image from one well, one field, one x,y-plane and one channel. This means for one well we had 7,500 tiff files (25 fields x 101 planes x 3 channels). We developed an automatic protocol that organized all tiff files from one well and saved them as zarr arrays. To do this, first we created an empty zarr array with size equal to a whole well image (the size of the image is collected directly from xml file). Then we put individual planes and channels from the same field together in the zarr array. Lastly, we stitched all the fields of a well together.

### 3D organoid segmentation

Organoid segmentation had two major steps, preprocessing of the image and segmentation of the organoids. Most of the algorithms used were from python scikit image processing package^20^. The preprocessing step reduces debris and intensity differences to increase accuracy of organoid segmentation. To do this: 1. The image is converted to gray scale, and a threshold is used to create a binary image, 2. Binary opening and dilation algorithms are then used to close small holes and remove small dots. For simplicity this is done on the 2D “max-projection” image, corresponding to the projection of the image maximum among all z-stacks. 3. The binary image is then multiplied by the original image to produce a “cleaned” image. For the organoid segmentation step: 1. The “cleaned” image is converted into gray scale, and a multidimensional gaussian filter is used to remove noise. 2. The threshold triangle method is then used to generate a binary image. Note: All the above steps are done on single field instead of an entire well image because fields often vary in noise and intensity. 3. The individual fields are stitched together. Objects in the stitched image are given a label using the “measure label” algorithm, 4. Unwanted small objects are removed using the “morphology remove small objects” algorithm. “Measure regionprops table” algorithm is used to generate a table of measurements for each organoid containing: label, centroid, 3D bounding box (bbox), volume, volume (bbox, filled, and convex), major and minor axis length, euler number, extent, intensity (max, min, and mean), and inertia tensor eigvals. 5. The output is stored in a csv file with the file name containing the row and column number of that well.

### StarDist-3D training

To annotate ground truth images for training a StarDist-3D model, we followed guidelines from Stardist^38^ GitHub page. Ground truth images were annotated manually using Labkit^36^ a plugin in Fiji. Every nucleus was given a label. In total 36 organoids were labeled. 10, 12 and 14 organoids had cells which were fluorescently labeled with Hoechst, EGFP and mCherry respectively. Hoechst labeling was performed to diversify the training set. Among these 36 organoids, 24 had been treated with cisplatin. In total 3,862 nuclei were annotated. The data was augmented using elastic deformation, noise/intensity shift, and flip / 90-degree rotation. For training, we used parameters based on an example from the StarDist GitHub. The model configuration was: anisotropy = (9.0, 1.0, 1.0), backbone = ResNet, number of rays = 64, patch size = (16, 128, 128), epochs = 400. The median object size was (2, 18, 18). Thus, a network view of (17, 30, 30) was used to make sure at least one nucleus is seen by the network. “KFold scikit learn model selection” was used to split data into six folds for cross validation, with 30 and 6 organoids for training and validation, respectively. The trained model was then used to segment nuclei in organoids.

### 3D nuclei segmentation

The segmentation of nuclei in organoids was done using the trained model. 3D bbox information of each organoid was used to extract each organoid in the saved zarr image of cells. To segment nuclei in the organoid: The organoid image was normalized. The trained model was used to segment individual nuclei, by giving each nucleus in the organoid a label. The “scikit image measure regionprops table” algorithm was used to generate a table of measurements for each nucleus: label, centroid, 3D bounding box (bbox), volume, volume (bbox, filled, and convex), major and minor axis length, euler number, extent, intensity (max, min, and mean), inertia tensor eigvals, and eccentricity (which was calculated as minor axis length/major axis length). Each fluorescence channel was processed and segmented separately. The data for all nuclei in a well were then stored as a csv file (one csv file per well) with the file name contains the row and column number of that well.

### Calculating area of imaged well with organoids

*Cellos* includes a customized method for distinguishing image areas that contain organoids in focus within the acquired image of every well (extended fig.7a-c). To do this, the whole well image with segmented organoids is used as input. To calculate the area all the processes are done on the 2D “max-projection” image. First, any very big objects that are not organoids are removed (note: we rarely have these objects), then any space between adjacent organoids is closed using “scikit image dilation and binary closing”. Second, the non-zero pixels (pixels with organoids) are counted using “Opencv-python (cv2) countNonZero” package. Finally, the number of pixels is converted into desirable unit like mm^2^, based on the size of an image pixel in μm. Notably, this step to calculate the area is done during organoid segmentation and can be skipped if not needed.

### Post-segmentation Analysis

The post-segmentation analysis used the csv files generated in the organoid and nuclei segmentation steps. Below are the analyses that were performed:

### IC_50_ estimation

Cell density for each well was calculated by dividing the number of segmented cells by the customized calculated area of the imaged well with organoids. Cell densities for each drug-treated well were compared to cell densities from control wells (medium only). IC_50_ values were estimated using R package *nplr (*N-Parameter Logistic Regression. R package). For the luminescence based IC_50_ assay, luminescence values were compared to control wells and IC_50_ values were calculated as mentioned above. Error bars represent the standard deviation of triplicates for each condition.

### Cells co-localization analysis

The co-localization analysis was performed on organoids consisting of two cell populations each labeled with either EGFP or mCherry. To calculate co-localization score of a specific fluorescently labeled clone, for example B-mCherry in an organoid: 1. Centroid of nucleus was computed for each cell in the organoid. 2. All pairwise distances (Euclidean distance) between nuclei centers were computed. 3. For each B-mCherry cell, the neighboring cells were ranked by distance in the window of K=5, 10, 20, 30, 40, and 50 cells. 4. B-mCherry cells proportion was then calculated in each window (K) and normalized for the overall proportion of B-mCherry cells in the organoid. This was performed to avoid the bias of varying proportions of B-mCherry cells across all organoids. 5. Co-localization score for B-mCherry in the organoid for a given K was then calculated by averaging the co-localization score of all B-mCherry cells in the organoid. We hereafter denote the co-localization score for clone type X in an organoid, *Lo* as,

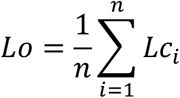

where, *n* is the total number of cells of clone type X in the organoid and *Lc* is the co-localization score of a cell of clone type X in a given K and is defined as,

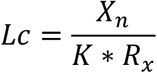

where, *X*_*n*_ is the number of cells of clone X in given K, and *R*_*x*_ is the proportion of clone X in the organoid.

### Statistical Tests

Details on statistical tests are in figure legends. All t-tests were two-tailed. All error bars indicate standard deviation. In all boxplots, the center line represents the median, the box limits represent the upper and lower quartiles, and the whiskers represent the 1.5x interquartile range.

## Data and code availability

Due to the size of data, we are providing raw images from one well along with its metadata as an example to run *Cellos* pipeline which can be found under: https://figshare.com/articles/dataset/cellos_data_zip/21992234. We are also sharing our entire *Cellos* pipeline and manually annotated training dataset which can be found on our GitHub page: https://github.com/TheJacksonLaboratory/Cellos. Other imaging data are available upon request.

## Acknowledgement

The authors gratefully acknowledge the contribution of the Flow Cytometry service, Single Cell Biology service, Cyberinfrastructure high performance computing resources, and Research IT at The Jackson Laboratory for expert assistance with the work described in this publication. The authors would like to thank Ali Foroughi Pour for helpful comments. The authors also would also like to thank Jim Peterson, Yi Li, Philipp Henrich, and Erick Ratamero for helpful discussions about organoids and nuclei segmentation. Research reported in this publication was supported by NIH grant R01CA230031 and NIH/NCI grant P30 CA034196.

## Author contributions

PM designed the study, generated training dataset, developed the software, performed data analysis, and wrote the paper. PK designed the study, generated training dataset, performed data analysis, generated experimental data, and wrote the paper. DM contributed to software development and reviewed the paper. SN performed data analysis and reviewed the paper. JN contributed to design of the study and reviewed the paper. MB and OA contributed to experimental design and reviewed the paper. EL designed the study, interpreted data, wrote the paper, and supervised the project. JC designed the study, interpreted data, wrote the paper, and supervised the project.

## Extended data

**Extended data fig 1:**
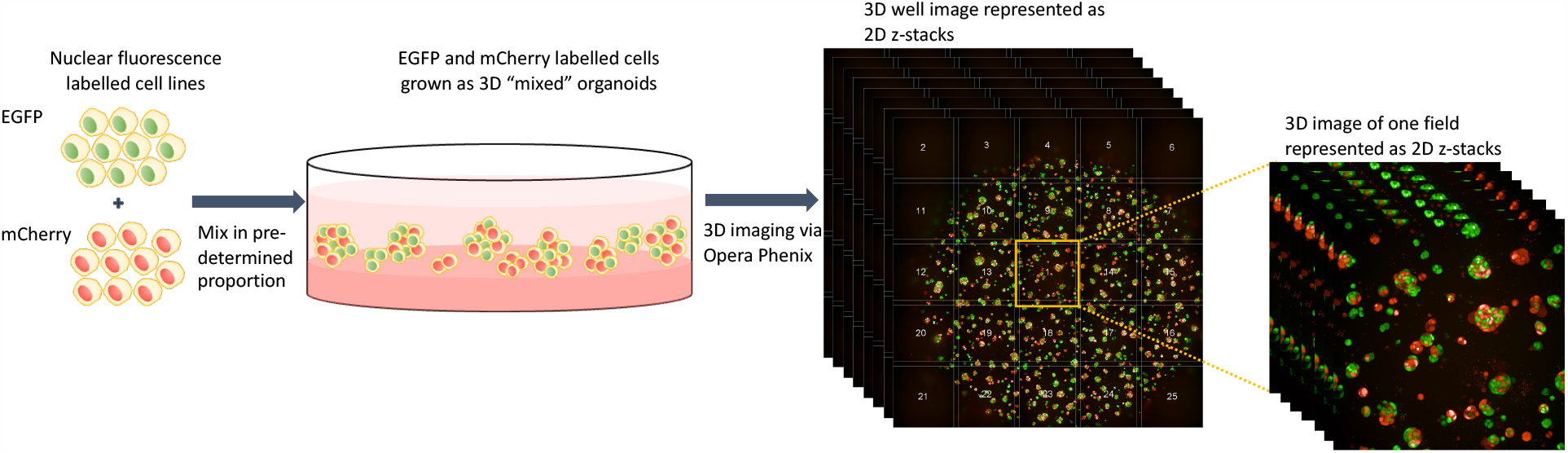
Schematic representation of 3D organoid culture and imaging platform. EGFP and mCherry labeled cells are mixed in pre-determined proportion in culture to form 3D “mixed” organoids. The Opera Phenix system is then used to image the organoids in 3D. 25 fields are imaged per well.

**Extended data fig.2:**
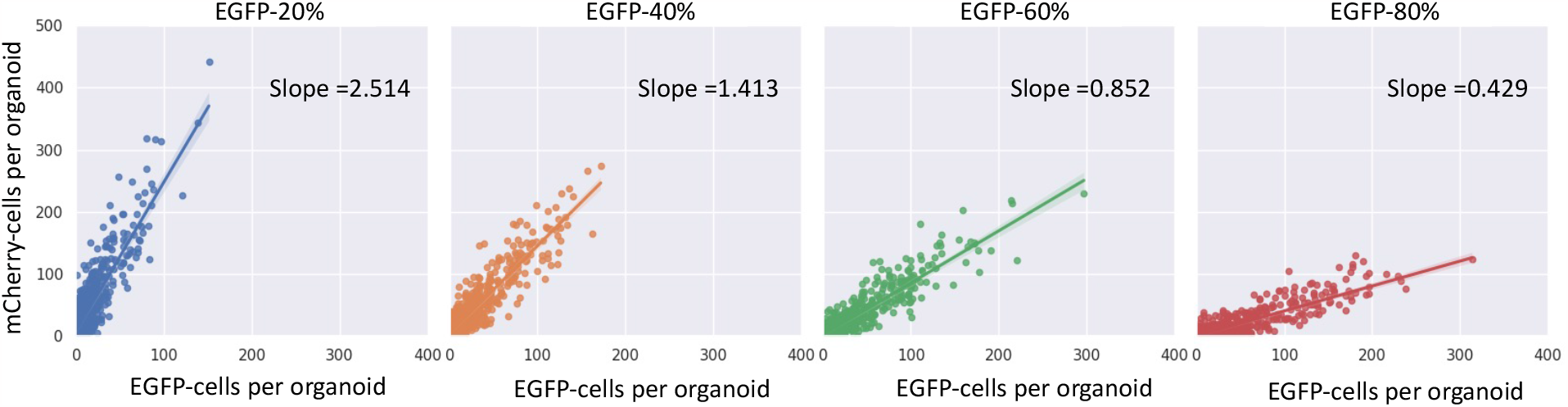
Quantification of fluorescently labelled cell populations at the organoid resolution. Number of EGFP vs mCherry cells detected in each homogeneously mixed B organoid. Each dot depicts an organoid. Seeding conditions of EGFP-20% (blue), EGFP-40% (orange), EGFP-60% (green) and EGFP-80% (red) are shown from left to right. Slope of the fitted linear regression is noted.

**Extended data fig.3:**
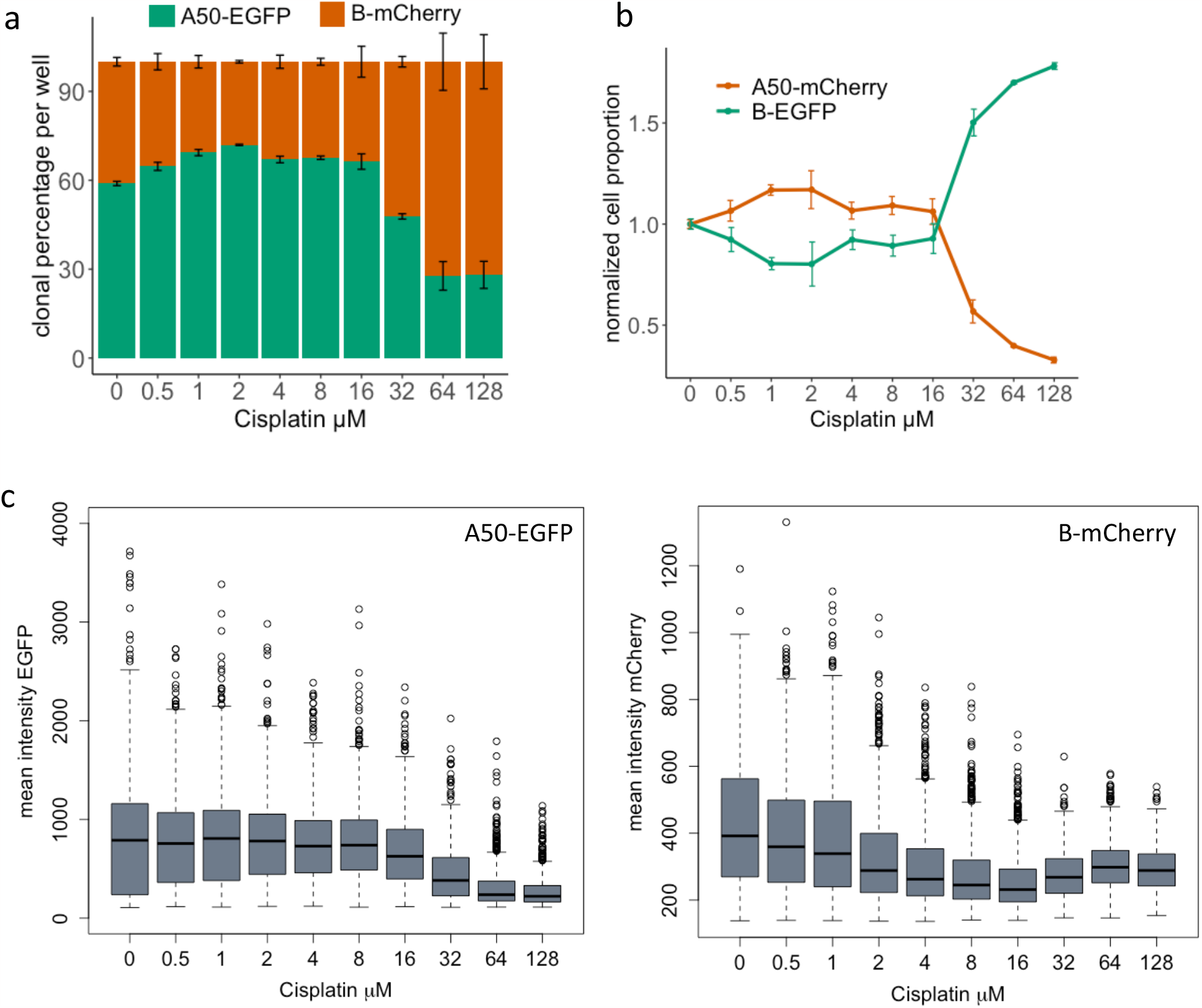
Quantification of clonal response to cisplatin treatment. **a**. Percentage of A50-EGFP and B-mCherry cells detected by *Cellos* for increasing concentrations of cisplatin treatment. Each well was treated for 4 days. Values represent mean clonal percentages across three replicates and error bars show standard deviation. **b**. Normalized cell proportions for A50-mCherry and B-EGFP clones when co-cultured as heterogeneously mixed organoids and treated with cisplatin for 4 days. Mean and standard deviation values of replicates for each condition are plotted. **c**. Mean intensity of EGFP (left panel) or mCherry (right panel) in heterogeneously mixed organoids consisting of A50-EGFP and B-mCherry. Organoids from three replicate wells are combined.

**Extended data fig.4.**
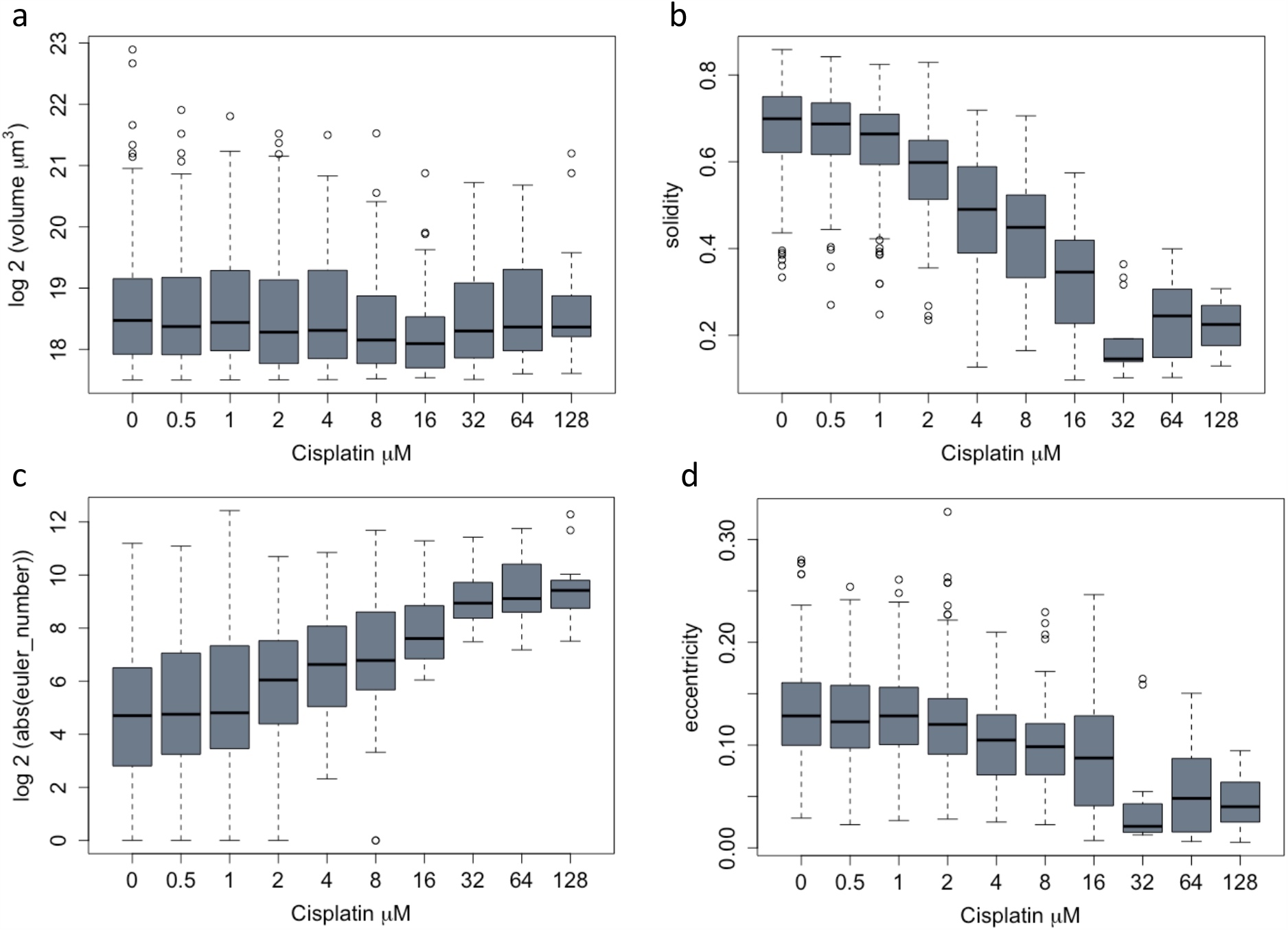
Changes in organoid morphologies due to cisplatin treatment. Volume (**a**), solidity (**b**), Euler number (**c**) and eccentricity (**d**) for segmented large organoids after exposure to range of cisplatin treatment for 4 days.

**Extended data fig.5:**
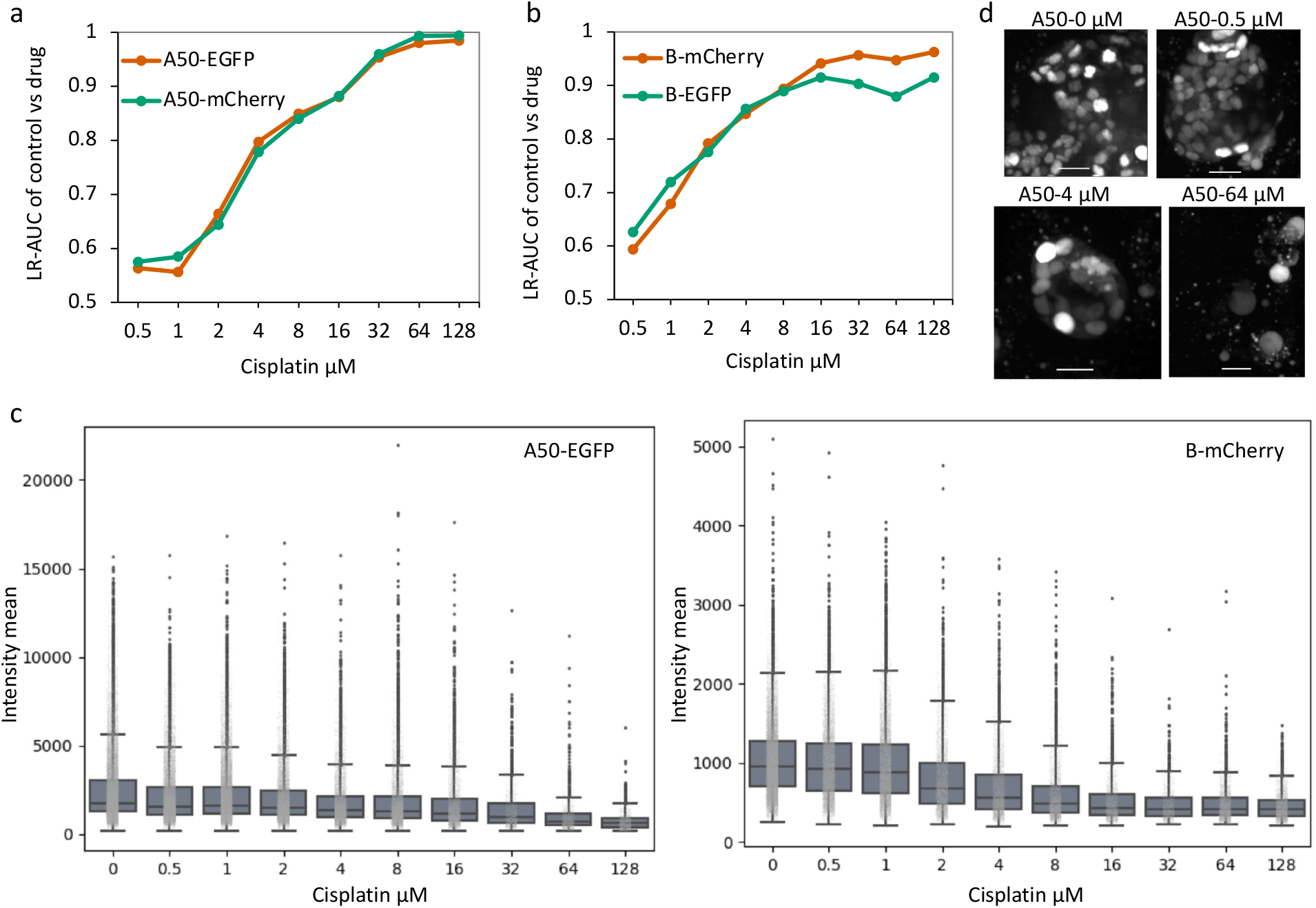
Changes in nuclear morphologies due to cisplatin treatment. **a**. Logistic regression classifications AUC of nuclear morphologies of A50-EGFP or A50-mCherry cells when comparing nuclei in control with nuclei exposed to cisplatin for 4 days. **b**. Logistic regression classifications AUC of nuclear morphologies of B-EGFP or B-mCherry cells when comparing nuclei in control with nuclei exposed to cisplatin for 4 days. **c**. Mean intensity of A50-EGFP (left) and B-mCherry (right) nuclei after cisplatin exposure for 4 days. **d**. Representative images of A50- EGFP cell nuclei at day 4 post select concentrations of cisplatin exposure. Scale bar represents 25 μm.

**Extended data fig.6:**
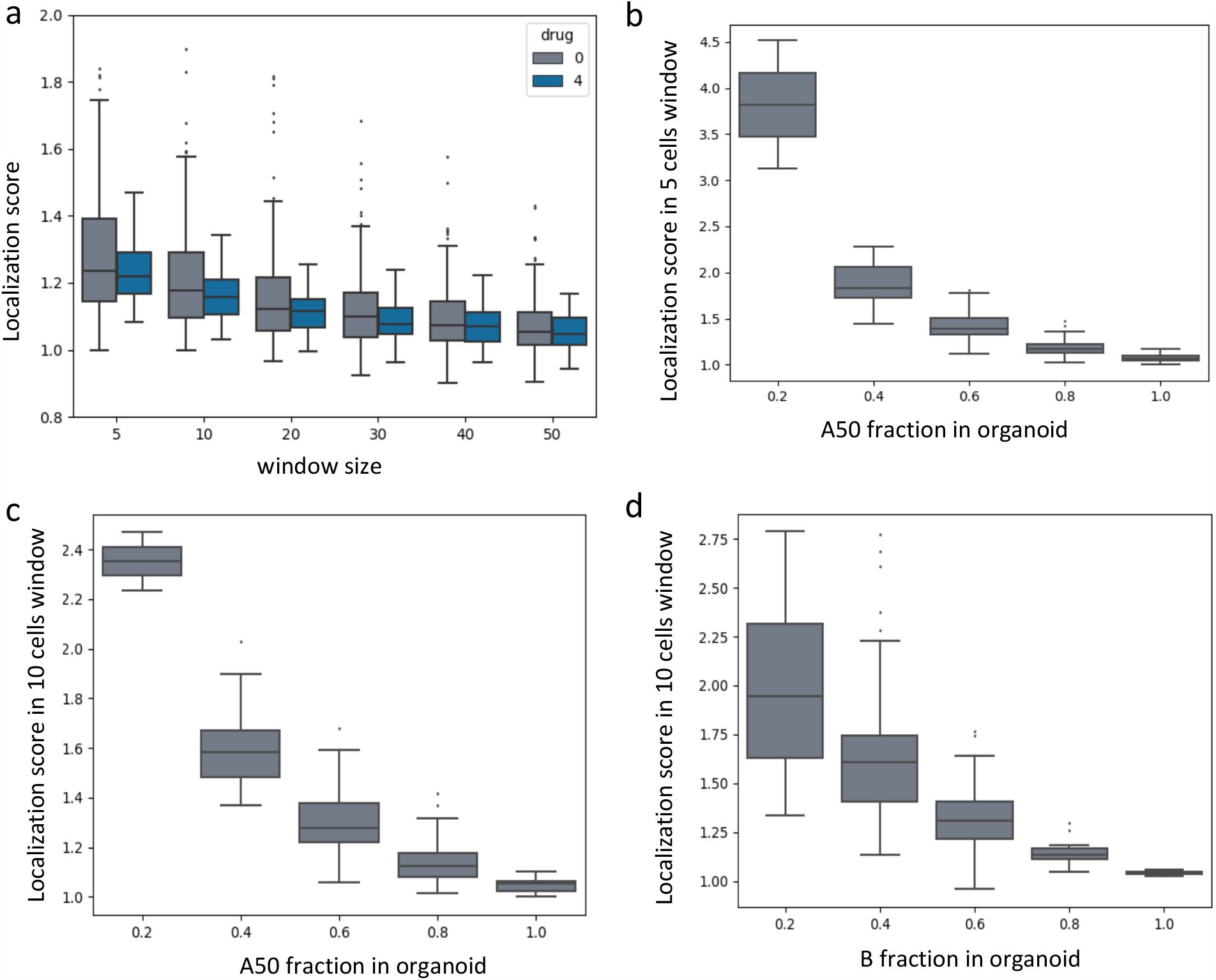
Identification of cellular localization in organoids by *Cellos*. **a**. Distribution of localization score vs. window size (5, 10, 20, 30, 40, 50) for A50-EGFP cells within heterogeneously mixed organoids. Organoids from control and 4 μM cisplatin exposure for 4 days were used for this analysis. **b, c** Localization scores for A50-EGFP cells in the 5 cell and 10 cell window size in organoids with varying proportions of A50 clone fraction are plotted. Organoids are grouped in bins with increments of 0.2 for the A50 cell fraction. **d**. Localization scores for B-mCherry cells (10 cell window size) in organoids with varying proportions of B clone fraction are plotted.

**Extended data fig.7:**
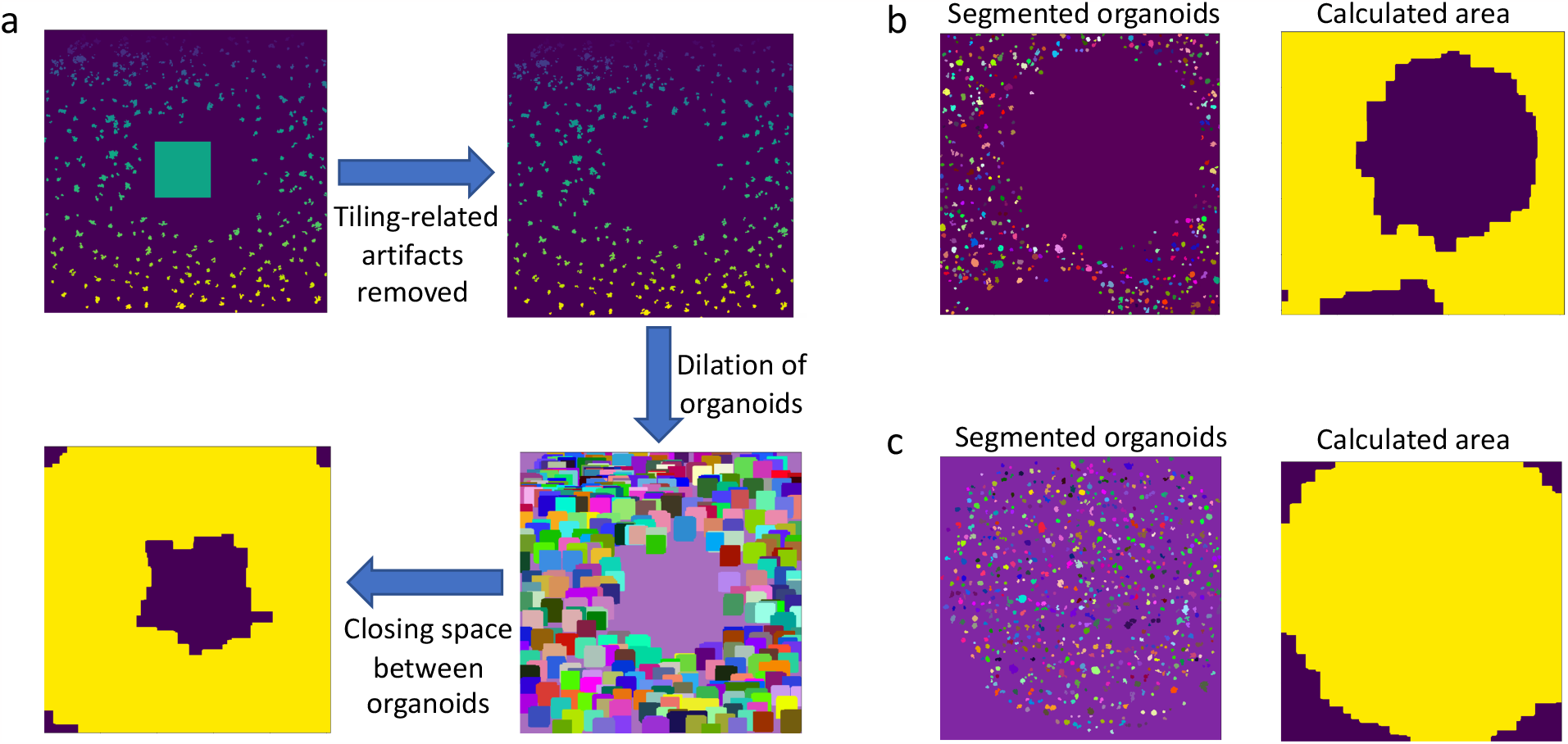
Computational process of calculating image area that contains organoids. **a**. Schematic representation of the pipeline to detect the area of the image with organoids in focus. **b, c**. Examples of segmented organoids image (left panel) and *Cellos-*detected area of the image with organoids in focus, shown in yellow (right panel). Images are z-axis maximum projections.

**Extended data table 1:** Detection of multiple fluorescently labelled cell populations in homogeneously mixed organoids. EGFP and mCherry labeled cells were mixed to form homogeneously mixed organoids of A50 or B clones. Pre-determined EGFP percentages of 20, 40, 60 and 80 were used and organoids were imaged at day 0 and day 4. Table showing percentage of EGFP labeled cells, and standard deviation (STDV) of three replicate wells for each condition

|  | Day 0 |  |  |  | Day 4 |  |  |  |
| --- | --- | --- | --- | --- | --- | --- | --- | --- |
|  | A50 |  | B |  | A50 |  | B |  |
| Seeded<br>EGFP(%) | EGFP(%) | STDV | EGFP(%) | STDV | EGFP(%) | STDV | EGFP(%) | STDV |
| 20 | 25.251 | 0.456 | 21.255 | 0.982 | 24.867 | 0.4 | 22.41 | 0.426 |
| 40 | 42.556 | 1.167 | 38.366 | 1.493 | 40.608 | 0.852 | 37.535 | 0.915 |
| 60 | 59.928 | 0.494 | 55.458 | 1.143 | 54.924 | 0.845 | 52.194 | 1.348 |
| 80 | 76.829 | 0.376 | 74.597 | 0.489 | 71.054 | 0.145 | 70.138 | 1.392 |

